# NestedBD: Bayesian Inference of Phylogenetic Trees From Single-Cell DNA Copy Number Profile Data Under a Birth-Death Model

**DOI:** 10.1101/2022.01.16.476510

**Authors:** Yushu Liu, Mohammadamin Edrisi, Huw A. Ogilvie, Luay Nakhleh

## Abstract

Copy number aberrations (CNAs) are ubiquitous in many types of cancer. Inferring CNAs from cancer genomic data could help shed light on the initiation, progression, and potential treatment of cancer. While such data have traditionally been available via “bulk sequencing”, the more recently introduced techniques for single-cell DNA sequencing (scDNAseq) provide the type of data that makes CNA inference possible at the single-cell resolution.

In this paper, we introduce a new birth-death evolutionary model of CNAs as well as a Bayesian method, NestedBD, for the inference of evolutionary trees (topologies and branch lengths with relative mutation rates) from single-cell data under this model. We assessed the accuracy of our method on both simulated and biological data and compared it to the accuracy of two standard phylogenetic tools, namely neighbor-joining and maximum parsimony (MP). We show through simulations that our method infers more accurate topologies and branch lengths. We also studied the ancestral state reconstruction accuracy with the birth-death evolutionary model and found it outperformed MP. Finally, running all three methods on a colorectal cancer data set, we observed that among all three methods, only the phylogeny inferred by NestedBD clearly separated the primary tumor cells from the metastatic ones, providing a more plausible history of the tumor cells.

## Introduction

Copy number aberrations, or CNAs, are somatic mutations that delete or amplify genomic regions and could cause cancer by amplifying oncogenes [1, 8] or deleting of tumor suppressor genes [27, 20]. CNAs are distinguished from copy number variations, or CNVs, which are typically germline mutations that serve as markers for population or evolutionary genetic studies. CNAs can vary in terms of the size of the genomic region that is amplified or deleted, the number of such events across the genome, as well as the rate at which they occur [21]. In particular, a CNA could amplify an entire genome or delete/amplify an entire chromosome [3, 31]. However, CNAs are often smaller, spanning thousands or fewer base pairs [34].

The accumulation of CNAs during cancer development and progression results in intra-tumor heterogeneity (ITH), where distinct CNA signatures characterize different groups of cells [7]. Elucidating ITH from genomic data is important for the diagnosis, prognosis, and treatment of cancer [16]. Single-cell DNA sequencing (scDNAseq) is ideal for inferring CNAs and ITH as it generates DNA sequence data from individual cells that are readily available for comparative genomic and evolutionary analyses [23]. Indeed, several methods have been developed for inferring copy number profiles from scDNAseq data [18], though their accuracy needs improvement [17].

In this work, we target the problem of inferring the evolutionary history of a set of individual cells using scDNAseq data, where each cell is defined by its copy number profile. That is, we assume the copy number profiles have been estimated already, and treat these as the input. Furthermore, we focus on focal CNAs that impact sub-chromosomal genomic regions, rather than whole genomes or chromosomes.

SCICoNE [15] and CONET [19] are two recent tools for simultaneous CNA detection and evolutionary history reconstruction on scDNAseq, leveraging the shared evolutionary history among single cells to infer CNAs. In this regard, both SCICoNE and CONET estimate a mutation tree, where a path from the root to a leaf defines the CNA signature of all cells attached to that leaf. The problem we focus on here, instead, is inferring a phylogenetic tree with branch lengths, with the two main goals of our work being to study the appropriateness of (1) an independent-bins assumption in these analyses and (2) a birth-death model of CNAs under this assumption. In studies of CNAs, it is common to partition the genome into bins, where each bin is a fixed number of nucleotides, rather than conduct the analysis at the resolution of individual nucleotides [18]. Given that CNAs naturally span many bins and CNAs could overlap over time, copy numbers in adjacent bins are *not* independent. Trying to model CNAs as events while taking into account such dependencies could result in intractable inference problems. Indeed, the MEDICC model developed by Schwarz *et al*. [30] aims to capture these dependencies, but inference under this model is very limited in terms of the size of the data given the prohibitive computational requirements [10]. Even if it is violated in practice, assuming independence among sites and loci has long been central to phylogenetic and phylogenomic inference formulations, mainly because it allows for much more efficient inferences. Here, we study the impact of assuming that copy numbers across bins are independent on the quality of phylogenetic inference. Furthermore, we propose the first formulation and inference method for copy number profile data from scDNAseq based on a birth-death model of copy numbers. We developed a new method, NestedBD, for Bayesian inference of phylogenetic trees from scDNAseq data under a birth-death model of copy number evolution assuming the bins are independent. The cells are also assumed to have been sampled at a single time point. NestedBD is implemented as a package in BEAST 2 [5], utilizing existing Markov chain Monte Carlo (MCMC) implementations and allowing for joint inference of trees and model parameters.

We assessed the performance of NestedBD on simulated and biological data and compared it to the performance of two commonly used methods, neighbor-joining (NJ) [28] and maximum parsimony (MP) as implemented in PAUP [32]. These two methods are readily applicable to CNA data since NJ requires pairwise distances among cells, which can be computed from the copy number profiles, and MP works directly on the copy number profiles and seeks a tree that minimizes the total number of copy number changes along its branches. Furthermore, these two methods are run in a way that assumes independence among the bins. We found that NestedBD infers more accurate tree topologies than the other two methods on the simulated data. It also provides accurate estimates of branch lengths. NJ, on the other hand, provides unreliable branch lengths (some of them are even negative), and MP does not estimate branch lengths (it can, of course, be used to estimate the minimum number of copy number changes on each branch of the tree, but these quantities differ from true branch lengths). Applying all three methods to a colorectal cancer data sample from [4], NestedBD obtained a more plausible evolutionary history.

Both the simulator and NestedBD are available at https://github.com/Androstane/.

## Methods

In this work, we assume that the genomes under consideration are partitioned into bins such that all genomes have the same number and sizes of bins. The copy number profile of a cell at each bin is an element of {0, 1, 2, …}. In practice, copy number profiles are estimated from scDNAseq data and consequently have errors in them [17]. In this paper, we do not consider the issue of error in the data. In the simulated data, this is easily accomplished since the true copy number profiles are available. For the biological data set, we used the estimated copy number profiles as is. Accounting for errors in the profiles is an important direction for future research.

### A birth-death evolutionary model of CNAs

To compute the likelihood of phylogeny, we first need an evolutionary model that defines the transition probability between copy number states. We model the copy number amplification and deletion by a constant rate birth-death process {*Z*(*t*), *t* ≥ 0} with state space *S* = {0, 1, 2,....}. *Z*(*t*) gives the copy number state of a bin at time *t*. We assume each copy is amplified with birth rate *λ >* 0 and deleted with death rate *μ*> 0. The transition rate at time *t* with *Z*(*t*) = *m* equals *mλ* if *Z*(*t* + Δ*t*) = *m* + 1 and *mμ*if *Z*(*t* + Δ*t*) = *m* − 1. Note that when *Z*(*t*) = 0, the transition rate becomes zero, suggesting neither amplification nor deletion from zero is allowed. Then we define the transition probability between copy number states as follows. Let *i* be the copy number state at the child node, *j* be the copy number state at the parent node, and *t* be the time between the parent node and the child node. Since there’s no prior information on the birth and death rates, we assume *λ* = *μ*= *r*. According to [12], the conditional probability P(*i*|*j, t*) is:

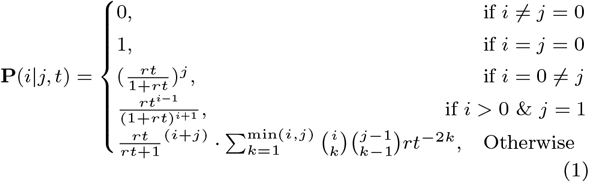

### Bayesian inference

Given the birth-death evolutionary model of copy number profiles, we use Markov chain Monte Carlo (MCMC) to sample from the following posterior distribution:

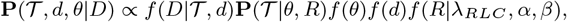

where *D* is the copy number profile, *θ* is the collection of parameters that define a birth-death^1^ tree prior on 𝒯, and *d* is the distance between the common ancestor of all cells and its diploid ancestor.

#### Prior

We assume the topology 𝒯 follows a two-parameter birth-death prior. Specifically, the birth-death model on the tree is a continuous-time process with two parameters, 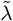 and 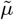, the instantaneous per-lineage rates of speciation and extinction, respectively, both of which are constant across the tree in their original characterizations [29, 24]. For the purpose of inference, we parametrize the model using the diversification rate 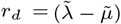 and extinction fraction 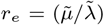, respectively. Since there is no prior information on the diversification rate and extinction fraction, we assume a uniform prior on both *r*_*d*_ and *r*_*e*_. In addition, the mutation rates on branches are assumed to follow the random local clock (RLC) model [9]. We assume a Poisson prior on number of rate changes with an expected value *λ*_*RLC*_ = log 2. This sets a 0.5 prior probability on the hypothesis of no change in mutation rate across the phylogeny. We also assume that rate multipliers are independently gamma distributed with *α* = 0.5 and *β* = 2 as in [9].

#### Likelihood

Assuming *r* = 1 with transition probability defined by birth-death evolutionary model on copy number state in Eq. (1), we used a modified Felsenstein’s pruning algorithm [11] to compute the likelihood of tree 𝒯 constructed from input copy number profile data *D*. We assume a diploid common ancestor of all tumor cells.

We define the state space of copy numbers as *S* = {0, 1, 2, …*k*}, where *k* ∈ ℕ defines the maximum copy number state to be considered during likelihood computation. For flexibility of the method, *k* is left to be a user-specified input with default being 9 considering the maximum value commonly observed in copy number state of cancerous cells. Note that although the likelihood computed under a larger *k* could be more accurate, it may not always be desirable computationally given the likelihood computation is 𝒪 (*nk*^2^), where *n* is the number of leaves in the tree.

Then, to compute the likelihood of a topology we adopt Felsenstein’s pruning algorithm with slight modification when computing the likelihood at the root to account for the diploid origin. Specifically, the original Felsenstein’s pruning algorithm computes likelihood across the whole tree using conditional likelihoods for all possible states at the root of the tree by ℒ = Σ_*x*∈*S*_ *π*_*x*_ *−_root_(*x*), where *x* refers to the copy number state at root node of the tree and *π*_*x*_ refers to the corresponding prior probability of that copy number state at the root of the tree. Our algorithm computes instead ℒ = Σ_*x*∈*S*_ P(*x*|2, *d*)* ℒ_root_(*x*), where P(*x*|2, *d*) corresponds to the transition probability as defined in Eq. (1) and *d* represents the time between the diploid and common ancestor of all cancerous cells. By default, *d* is inferred jointly with the topology by assuming a uniform prior on it.

### Evaluating tree inference on simulated data

#### Simulation protocol

To simulate data, we made three modifications to the CNA evolutionary simulator described in [17]. (1) The simulator now accepts a user-defined topology and simulates CNAs along the branches of the input tree. (2) If an input tree is specified by the user, the number of CNAs on a branch with length *t* is sampled from *Poisson*(*c · t*), where *c* is a user-specified parameter that controls the number of CNAs at the leaves of the tree. (3) The chromosome and position of the allele where a CNA occurs are sampled from a random distribution whose probability density function can be expressed as 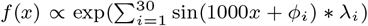 with *x* representing the location on genome. We fixed *ϕ*_*i*_ ∼ *Uniform*(−*π, π*) and *λ*_*i*_ ∼ *Uniform*(0, *a*), where *a* is a user-specified parameter that controls the non-uniformity of the distribution. When setting *a* = 0, the position of the CNA is sampled from a uniform distribution as in the original simulator.

To simulate CNAs with a similar pattern to the original CNAs in real data set, we used the maximum clade credibility (MCC) tree inferred by NestedBD on the biological data set as input to the simulator. The branch lengths of the MCC tree are summarized using median node heights across all sampled trees. CNAs are added along the branches of the tree with the number of CNAs sampled from a Poisson distribution whose mean is proportional to branch length. Each node has its unique CNAs on the corresponding branches while inheriting the CNAs from their ancestors. The copy number profiles of the leaves are then inferred using Ginkgo [14] with binning option of `variable _175000 _48 _bwa`. For the purpose of this study we used *c* = 125, *a* = 0.6, *X* = 1, *m* = 2Mbp, *e* = 100Mbp, with all other parameters set to their default values. We also assume there is no whole-genome amplification or deletion. We found the CNAs simulated under such conditions best resemble what we observed from the biological data set. On average, more than 90% of the CNAs overlapped with at least one other CNA. Finally, to satisfy the independent-bins assumption, we sample the bins with a 1*/*20 sampling rate before making the copy number profile data available to the inference methods.

#### Inference methods

For each simulated data set, NestedBD is run using the BEAST 2 implementation with coupled-MCMC [2] for 80 million iterations. Five chains with random seeds are run to assess the convergence of the MCMC runs. To summarize the posterior distribution, 2000 samples are taken from the MCMC chain for computation of inferred topology and branch length. We obtained 100 bootstrap replicates for each data set using both MP and NJ with PAUP [32]. Both MP and NJ return unrooted trees by default while NestedBD infers a rooted tree by assuming a diploid origin. To root the MP and NJ trees, we added as an outgroup a diploid “genome” and rooted the inferred tree at it.

#### Assessing the error rates of methods

We compute the true positive rates of reconstructed branches considering all phylogenies sampled from the posterior distribution of NestedBD or the bootstrap samples of MP and NJ for all simulated repetitions. More specifically, for each branch in the true tree, we count the proportion of trees in the posterior distribution or bootstrap samples having that branch.

We assess the the ability of a method to recover a branch in the true tree as a function of the length of that branch. To make sure the slopes across different methods are comparable, we fitted an ordinary least squares (OLS) regression of log-scaled branch lengths with respect to the true positive rate assuming the slopes share a common intercept.

#### Tree scaling

Branch lengths inferred by NestedBD are not in the same unit as those in the true tree. Therefore, to assess the accuracy of branch length reconstruction, we needed to scale the inferred phylogeny before comparing branch lengths. Given an inferred tree 𝒯 and a true tree *R* with same set of leaves, we find the scale factor by computing an OLS regression as follows. Let the set of clades in 𝒯 be 𝒯_*C*_ and the set of clades in *R* be where Y is a vector of true node heights and X is a vector of *R*_*C*_. We compute *β* that minimizes residue *R*(*β*) = ||Y − *β*X||^2^, inferred node heights of clades in 𝒯_*C*_ ∩ *R*_*C*_.

To compute the 95% highest posterior density (HPD) intervals and *R*^2^ measures from posterior samples of NestedBD, we summarized the trees from the posterior distribution by MCC trees with median node heights. We then computed the scaling factor of the MCC tree with respect to the true tree for each simulated data set. The same scaling factor is used to scale all selected samples from the posterior distribution.

### Evaluating ancestral profile inference on simulated data

We used the same simulated data to evaluate the performance of ancestral profile inference. For each data set, we inferred ancestral profiles on both the true tree as well as the inferred trees (MP tree as inferred by maximum parsimony, and MCC tree as inferred by NestedBD). The former allows us to assess the accuracy of methods assuming the tree is correct, whereas the latter allows us to factor in the tree estimation error when computing the accuracy of ancestral profile reconstruction.

For MP ancestral profile reconstruction, we used PAUP [32]. For NestedBD ancestral profile reconstruction, we used a dynamic programming algorithm adopted from [25]. The algorithm maximizes the joint likelihood given the binned copy number profiles of the single cells and a tree topology. The transition probability is computed by Eq. (1). To account for the diploid origin of all cells, we compute the probability of each state at the root by P(*x*|2, *d*) as defined in Eq. (1).

Hamming distances between true and inferred ancestral profiles are used for measuring accuracy. In the case where the true tree was used to reconstruct ancestral profiles, there’s a one-to-one correspondence between ancestral profiles on both trees (as the trees are identical). When inferred trees are used to reconstruct ancestral profiles, such a correspondence might not exist since the true and inferred trees could differ. In this case, a node *u* in the true tree has a corresponding node *v* in the inferred tree if and only if the set of leaves under *u* and the set of leaves under *v* are identical. Hamming distances of ancestral profiles of only corresponding nodes were computed in this case.

## Results

### Performance on simulated data

We discuss here the accuracy of the methods in terms of inferring the trees—topologies and branch lengths—and ancestral copy number profiles.

#### Accuracy of inferred topologies

While it is common to calculate the Robinson-Foulds (RF) distance [26] between the inferred tree and true tree to quantify their difference, this is not particularly useful in our case for at least two reasons. First, there are several groups of cells where cells within each group are equidistant from each other, and their resolution in a binary tree in arbitrary. The RF distance would heavily penalize resolutions that differ from the true one. Second, in analyses of real scDNAseq data, full resolution of the tree down to the individual cells is not of interest; instead, biologists are interested in resolution down to the level of sub-clones only. Therefore, we focus on a method’s ability to reconstruct a branch in the true tree as a function of the length of that branch. More specifically, we computed the true positive rate for a given branch length as the number of branches of that length that are correctly inferred by the method of interest. Furthermore, we fit the true positive rate data of each method using linear regression to study how the accuracy of each method improves with increasing branch length. The results are shown in Fig. 1.

**Fig. 1:**
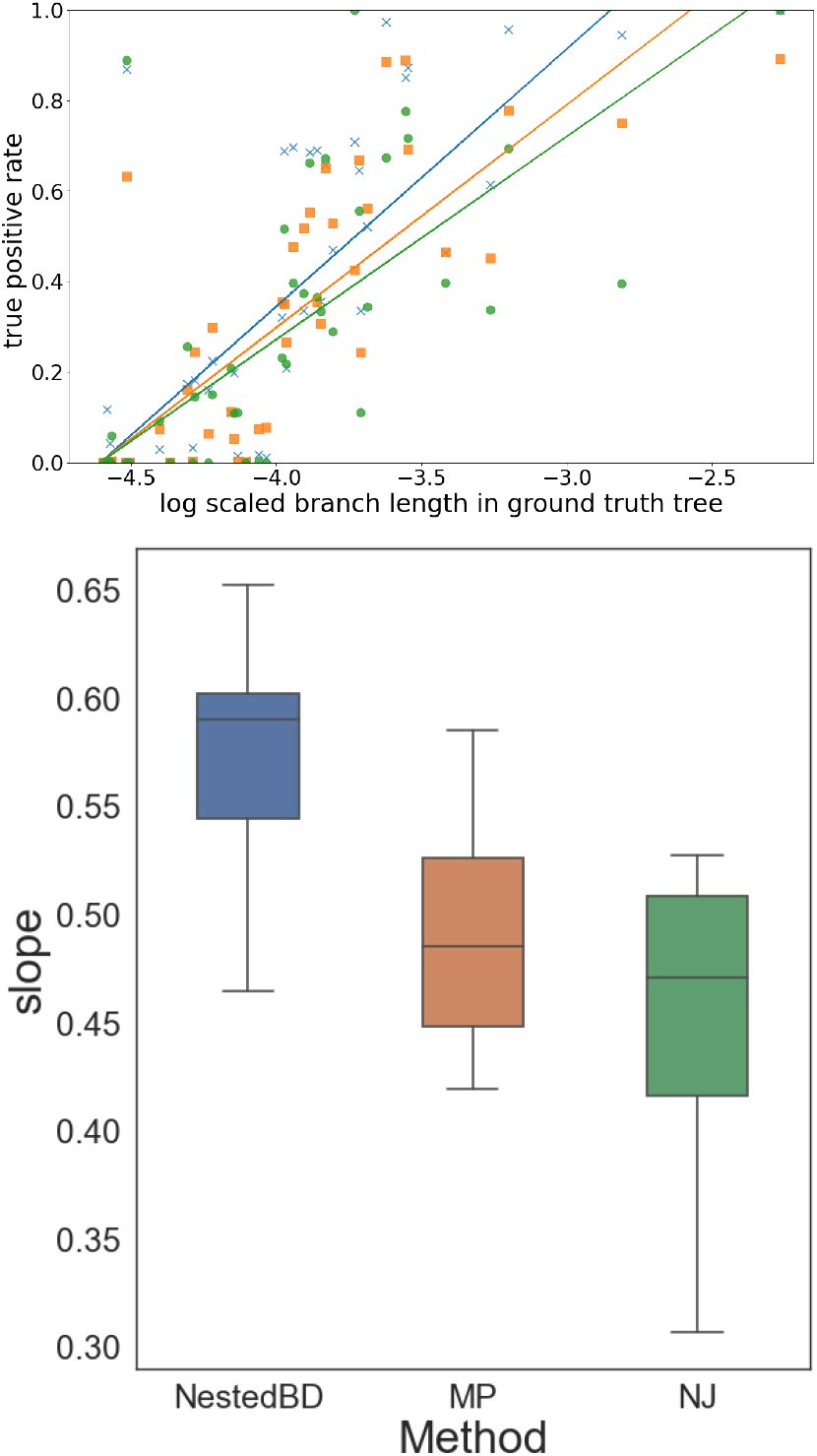
Accuracy of the summarized posterior distribution of reconstructed phylogeny. (Top) True positive rates of branches in the true tree vs. lengths of the branches. Each point corresponds to a branch length in the true tree (there are 10 true trees) and the proportion of trees (in the posterior or bootstrap samples) that have that branch, inferred by each of the three methods. ×: NestedBD; ▄: MP; •: NJ. A fitted OLS regression on all points for each method is shown. (Bottom) Box plots of the 10 slopes of linear regressions fitted for each method on the 10 simulated data sets. The whiskers correspond to the minimum and maximum values, while the line within the box corresponds to the median.

As expected, the longer a branch in the true tree, the better the method’s ability to recover it in the inferred tree. Furthermore, NestedBD has the best accuracy, followed by MP, followed by NJ. As the top panel of Fig. 1 shows, a large number of the true branches are very short and these tend to be closer to the leaves. As we discussed above, recovering branches near the leaves is not of much biological interest in studies of single-cell data.

#### Accuracy of estimated branch lengths

To the best of our knowledge, NestedBD is the first method that jointly infers branch lengths with mutation rates when reconstructing the evolutionary history of copy number profiles. MP constructs only the topology without branch length information, while NJ yields branch lengths that are often not biologically meaningful (the branch lengths might even be negative). We focus here on the accuracy of branch length estimates of NestedBD, and report them in terms of the node heights.

Since an inferred tree topology might differ from the true tree topology, we only compared the heights of nodes in the true tree that have corresponding nodes in the inferred tree. Furthermore, as described above, node heights were scaled to ensure comparability between the true and inferred node heights.

As in the case of topological accuracy, we evaluate the accuracy of estimated node heights using samples from the posterior distribution obtained by NestedBD. Fig. 2 summarizes the 95% HPD of node heights inferred by NestedBD after scaling (described in Section 2.3) for each of 10 simulated data sets. We also computed the *R*^2^ value of inferred and true node heights. As the figure shows, NestedBD obtains very accurate estimates of node heights. In particular, for all data sets, with the exception of data set I, the true node height is within the predicted 95% HPD interval for a majority of the nodes. Furthermore, for a large number of the nodes, the median of inferred node heights in the posterior sample appears to be a reasonable point estimate of the true node height.

**Fig. 2:**
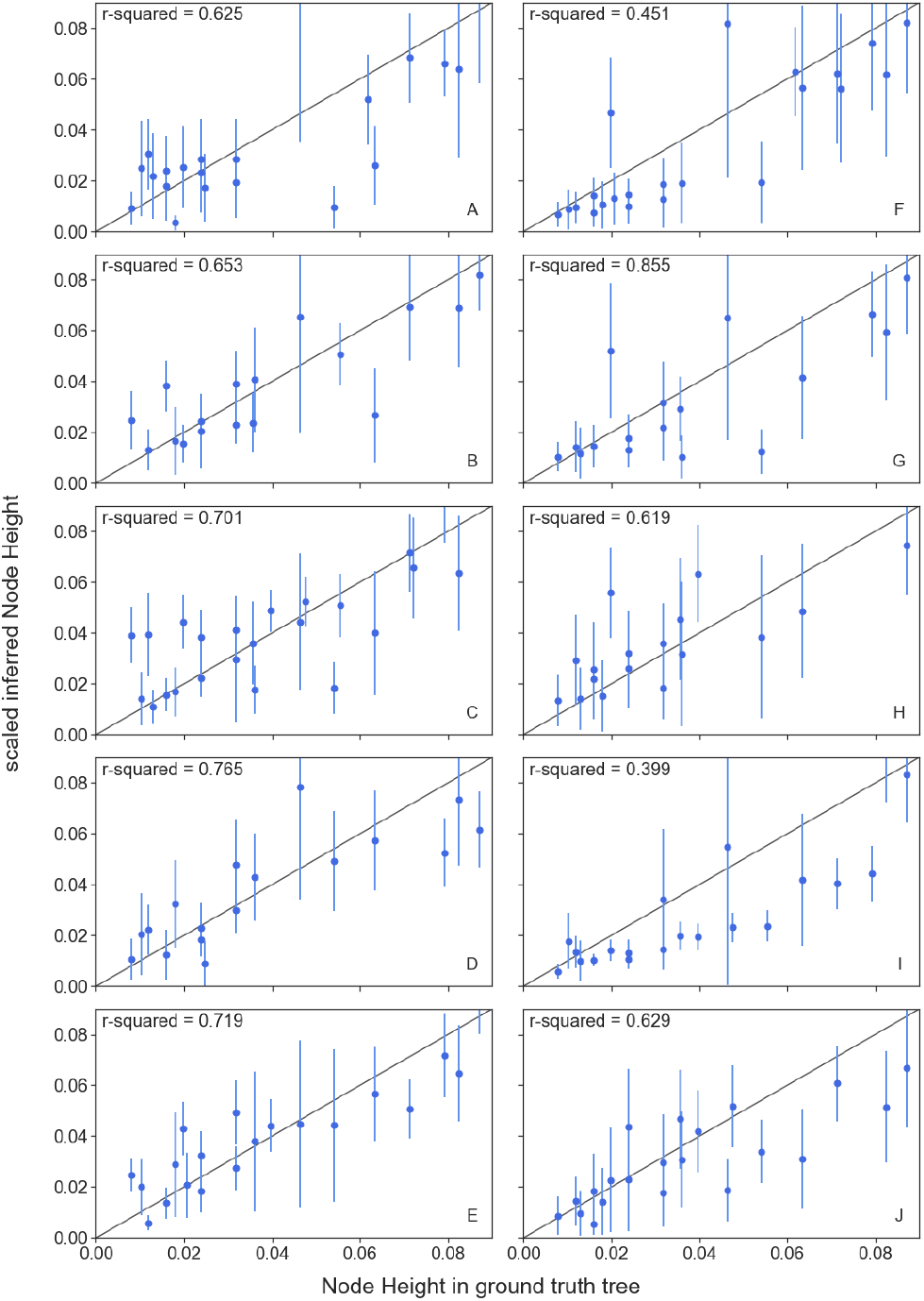
95% HPD of node heights estimated by NestedBD. Each panel corresponds to one of the ten data sets. For each data set, the node height inferred by NestedBD is summarized by both the median and 95% HPD interval of samples in the posterior distribution. The r-squared value (*R*^2^) is computed by fitting a linear model using the median of inferred node heights versus true node heights.

#### Accuracy of ancestral profile reconstruction

For ancestral state (copy number profile) reconstruction, we ran both NestedBD and MP on both the true trees and trees inferred by NestedBD. Furthermore, we estimated ancestral states using MP on the trees inferred by MP. NestedBD cannot be run on the trees inferred by MP since the tree does not have branch length estimates that are needed from NestedBD. Results are shown in Fig. 3.

**Fig. 3:**
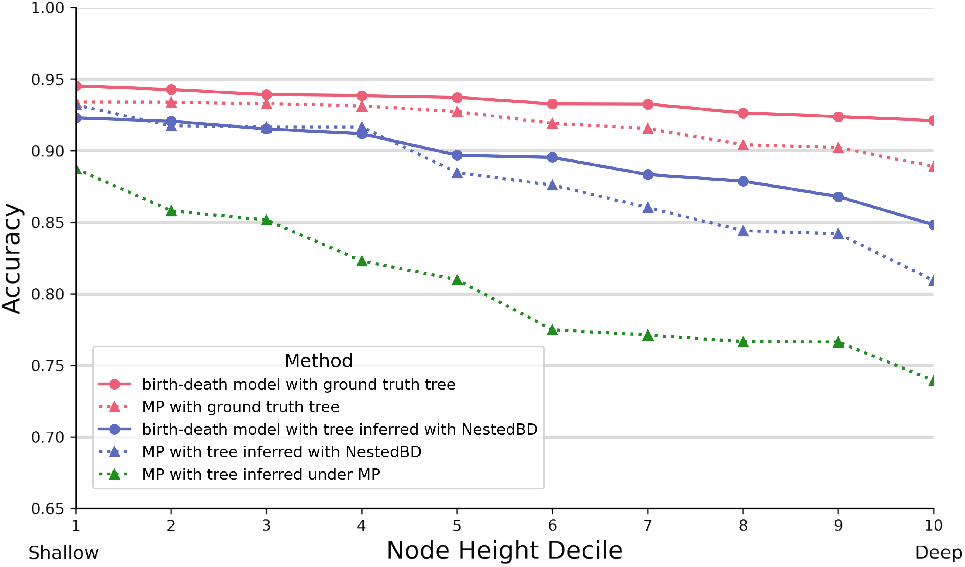
Accuracy of ancestral state reconstruction. True node height deciles in the true tree are shown on the x-axis.

A few observations are in order. First, in all scenarios, ancestral profiles are much more accurate at the shallower nodes (ones closer to the leaves) and that accuracy drops as the nodes become deeper. This might seem unintuitive given that the genomes at deeper nodes have fewer, and potentially less complex, CNAs with respect to the (normal) diploid genome. However, the data used for ancestral reconstruction is the profiles at the leaves of the tree, and the shallowest nodes are closest to those. Since reconstruction proceeds in bottom-up (leaves to root) fashion, errors in shallow reconstructed profiles could propagate to the deeper ones (given the dependencies of the latter on the former ones). Second, the quality of the tree has a big impact on the accuracy of ancestral profile reconstruction. This is unsurprising. The accuracy is highest when the true tree is used, followed by the cases where the tree inferred by NestedBD is used, and finally using the MP tree yields poorest results. As we showed above, trees obtained by NestedBD are more accurate than those obtained by MP. Third, while for the shallowest nodes, the accuracy of ancestral profile reconstruction by NestedBD is comparable to that of MP, the gap between the two methods opens up in favor of NestedBD as nodes become deeper. This, too, makes sense. As the node gets deeper in the tree, the likelihood that a bin in the genome undergoes multiple CNAs becomes higher (not only mathematically, but this has been hypothesized in the case of cancer genomes [6]). By definition of the parsimony criterion, MP is not well equipped to handle recurrent mutations at the same locus, whereas such mutations are accounted for in the birth-death process underlying NestedBD. Finally, as we discussed above, from a tree inference perspective, the shallowest nodes are harder to infer than deeper one. Since MP makes more errors in tree inference and those errors are mostly in shallow nodes, the performance of ancestral profile reconstruction by MP on the tree inferred by MP drops rapidly with the node height.

### Analysis of a colorectal cancer sample

We applied NestedBD to a single-cell copy number profile data set from colorectal cancer patient CRC04 obtained from [4]. We took a subset of the data set by randomly sampling 52 cells taken from the primary tumor site (PT) and Lymph node metastasis (LN) after excluding cells taken from normal adjacent tissue. For analysis of biological data set, NestedBD was run for 80 million iterations to ensure convergence of the MCMC chain. We then computed the MCC of the trees from the posterior samples, which is shown with a heat-map summarizing copy number profiles in Fig. 4. We inferred the ancestral profiles using NestedBD, and annotated the branches that define the primary tumor cell clades with colorectal-cancer-related genes, according to [33], that were impacted by CNAs. The inferred tree has a relatively long branch that separates the normal cell and the most recent common ancestor of all 52 tumor cells. This observation supports a punctuated mode of evolution of the tumor [13]. Nine colorectal-cancer-related mutations are detected on this branch, including in *APC*, a well-recognized initiator gene in colorectal cancer [22].

**Fig. 4:**
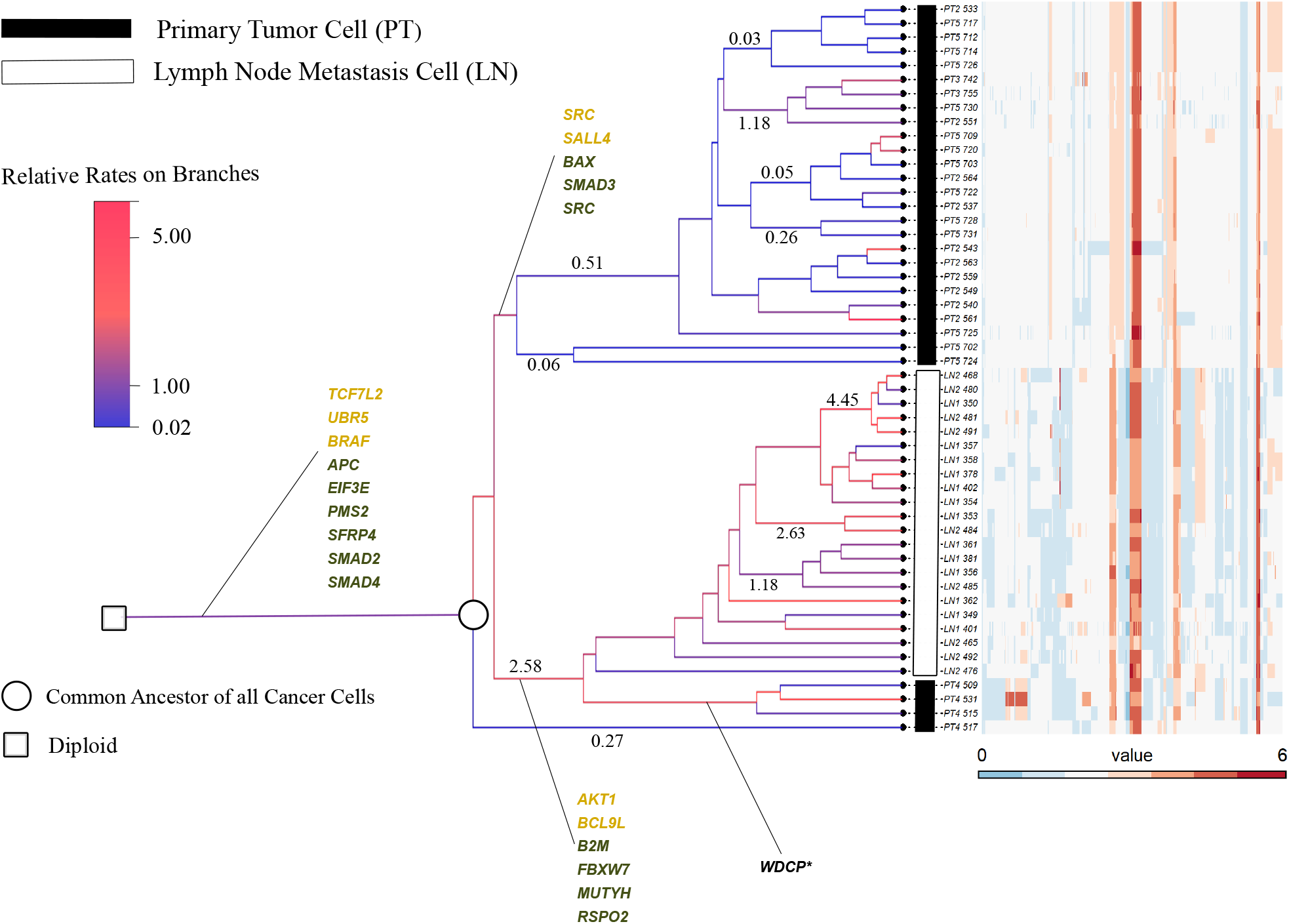
Inference results using NestedBD on data from colorectal cancer patient CRC04 from [4]. The heatmap shows the copy number profiles of the sampled cells. Colors of the branches indicate the estimated branch-specific relative rates from fast (red) to slow (blue). Branches defining the primary tumor cell clades are annotated with colorectal-cancer-related oncogenes (yellow) and tumor suppressor genes, or TSGs (green), impacted by CNAs. Note: while WDCP is netiher an oncogene nor a TSG, the WDCP protein has been identified in a fusion protein with ALK in colorectal cancer [35].

After that, a group of primary tumor cells acquired six additional mutations (in genes *AKT1, BCL9L, B2M, FBXW7, MUTYH, RSPO2*). An increase in mutation rate is also observed on this branch. Part of the cells then metastasized to the lymph nodes and evolved with a relatively high mutation rate; the rest remained at the primary tumor site (PT4). The rest of the cells at the primary tumor site acquired several unique mutations (in genes *SRC, SALL4, BAX, SMAD3, SRC*), but with a slower mutation rate (PT2 PT3, PT5). We also applied MP and NJ to the same data set; results are shown in Fig. 5. We observe that while MP places the same set of primary tumor cells (PT4) under the LN lineage, the topology seems to suggest those primary tumor cells are derived from the LN lineage, which is an unlikely evolutionary scenario. NJ infers a more reasonable topology similar to that inferred by NestedBD. While the branch lengths inferred by NJ are not biologically meaningful (especially considering that the inferred tree should be very close to ultrametric given that the cells were sampled at a single time point), we observed that the LN lineage is closer to the diploid cell at the root in the NJ tree. This observation is consistent with the higher relative mutation of the LN lineage in the tree inferred by NestedBD.

**Fig. 5:**
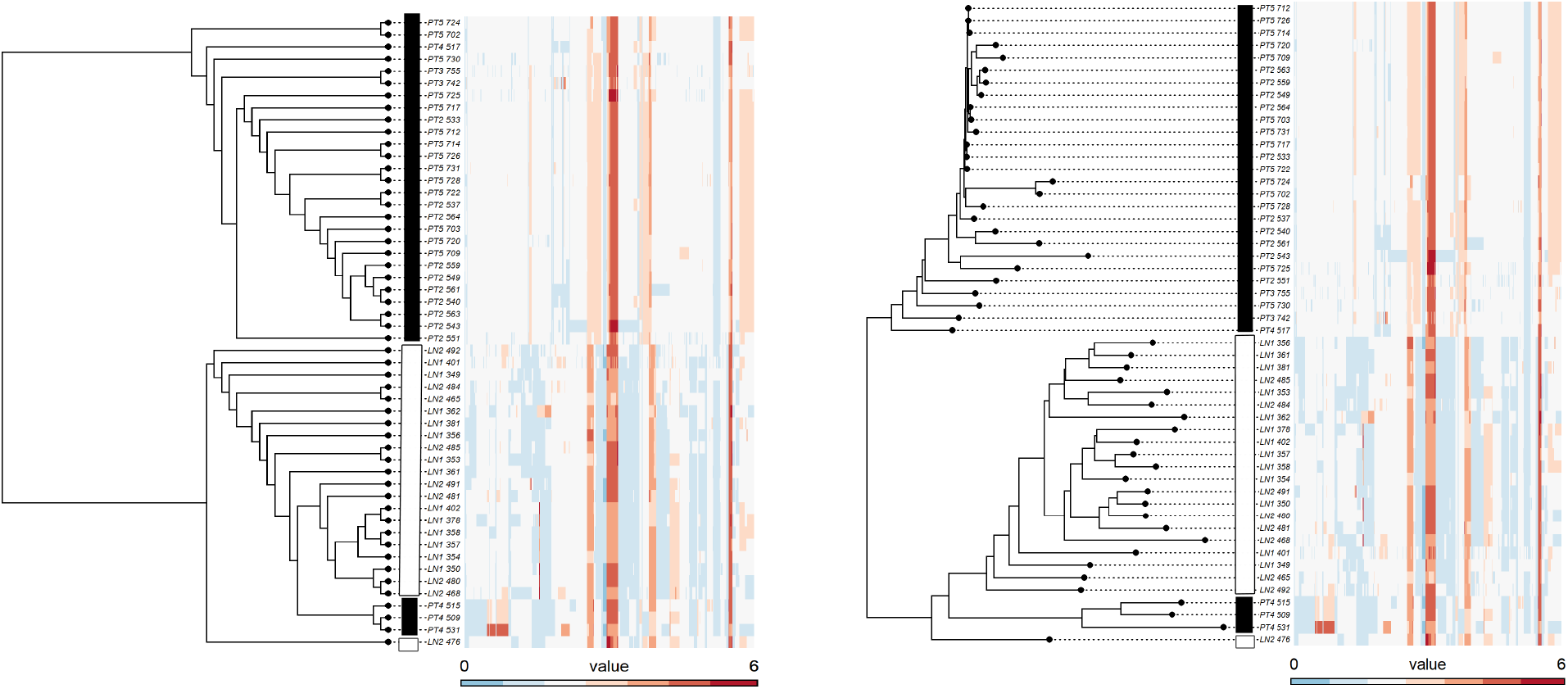
Inference results using MP and NJ on data from colorectal cancer patient CRC04 from [4]. The heatmaps show the copy number profiles of the sampled cells. (Left) Tree inferred by MP. (Right) Tree inferred by NJ. At the leaves of the trees, solid rectangles correspond to primary tumor cells, and open rectangles correspond to lymph node metastasis cells.

## Discussion

In this paper, we presented NestedBD, a Bayesian method for joint inference of evolutionary trees and branch lengths from scDNAseq copy number profiles. Specifically, we proposed a novel evolutionary model that uses a continuous birth-death process to model copy number amplification and deletion, accounting for the fact that there could be multiple CNAs at a single bin. We assume the phylogeny also follows a birth-death branching process parameterized by a diversification rate and an extinction fraction, and branch-specific mutation rates so that it is possible to distinguish between rapid expansion and slower mutations. NestedBD also infers the distribution of birth and death rates on the tree topology, the relative time between (normal) diploid cells and the most recent common ancestor of tumor cells. A major distinguishing feature of NestedBD is that it infers a tree with branch lengths representing the relative times of the tumor phylogeny nodes. NestedBD is implemented as a BEAST 2 package to utilize efficient implementation of MCMC. We assessed the accuracy of NestedBD on simulated data, demonstrated its application to a biological data set, and compared that to results obtained by two existing methods, namely maximum parsimony and neighbor-joining. NestedBD provides more accurate results overall.

To the best of our knowledge, NestedBD is the first method to infer a tree with branch lengths that measure relative times of evolution given single-cell copy number profiles (assuming independence among bins). While the simulated data do not assume independence among bins and biological data are very unlikely to satisfy such an assumption, the results we obtained demonstrate that utilizing the independence assumption for computational efficiency does not necessarily impact inference quality much. Recent methods, such as [19] and [15], infer breakpoints and clonal trees simultaneously, but they do not provide information on the times of nodes or mutations on the tree.

Two directions for future research are (1) accounting for error in the copy number profile estimates, since in practice these are estimated from genome read data, and (2) developing an inference method that works on the genome read data directly so that it simultaneously infer the copy number profiles and evolutionary history. While the latter is expected to produce the most accurate results, its scalability to large data sets could prove very challenging.

## Acknowledgments

This work was supported in part by National Science Foundation grants IIS-1812822 and IIS-2106837.

## Footnotes

1 There are two birth-death processes employed in this work— one on the shape of the trees and another on the copy number states. There are distinct and should not be confused.

